# *In vivo* emergence of high-level resistance during treatment reveals the first identified mechanism of amphotericin B resistance in *Candida auris*

**DOI:** 10.1101/2021.10.08.463721

**Authors:** Jeffrey M. Rybak, Katherine S. Barker, José F. Muñoz, Josie E. Parker, Suhail Ahmad, Eiman Mokaddas, Aneesa Abdullah, Rehab S. Elhagracy, Steve L. Kelly, Christina A. Cuomo, P. David Rogers

## Abstract

*Candida auris* has emerged as a healthcare-associated and multidrug-resistant fungal pathogen of great clinical concern. While as much as 50% of *C. auris* clinical isolates are reported to be resistant to amphotericin B, to date, no mechanisms contributing to this resistance have been identified. We report here mutations in the *C. auris* sterol-methyltransferase gene, *ERG6*, as the first identified mechanism of amphotericin B resistance in this emerging pathogen and describe the clinical case in which this high-level amphotericin B resistance was acquired *in vivo* during therapy. Whole genome sequencing revealed the four *C. auris* isolates obtained from this single patient case to be genetically related and identified a mutation in *ERG6* as being associated with amphotericin B resistance. Cas9-mediated genetic manipulations confirmed this mutation alone to confer a >32-fold increase in amphotericin B resistance, and comprehensive sterol profiling revealed an abrogation of ergosterol biosynthesis and a corresponding accumulation of cholesta-type sterols in isolates and strains harboring the clinically derived *ERG6* mutation. Together these findings represent the first significant step forward in the understanding of clinical amphotericin B resistance in *C. auris*.

## INTRODUCTION

In little over a decade, *Candida auris* has transformed from a newly identified species of *Candida* originally associated with infections of the auditory canal, to being recognized by the Centers for Disease Control and Prevention (CDC) as the first fungal pathogen to represent an urgent level of threat to public health.[1-4] Recently found to be the cause of outbreaks of invasive candidiasis on multiple continents and having been isolated from patients in more than 45 countries, *C. auris* readily colonizes patients and disseminates easily within healthcare facilities, changing the paradigm of healthcare-associated fungal infections.[4, 5] Moreover, *C. auris* frequently exhibits reduced susceptibility to multiple agents of the already limited antifungal armamentarium.[1, 4, 6] While limited epidemiological and clinical outcomes data currently preclude the establishment of true clinical breakpoints for the assessment of *C. auris* antifungal susceptibility, the CDC has defined tentative antifungal breakpoints using susceptibility data from hundreds of clinical *C. auris* isolates and available pharmacokinetic-pharmacodynamic data. Applying these tentative breakpoints, more than 90% of *C. auris* isolates are resistant to fluconazole (modal minimum inhibitory concentration [MIC] ≥256 mg/L), 30 to 50% are resistant to amphotericin B, and approximately 5% are resistant to echinocandins.[4, 7] Unfortunately, *C. auris* has also demonstrated the capacity to rapidly acquire resistance to antifungals *in vivo*, leaving clinicians with no reliable option for the treatment of infections caused by this emerging public health threat.[8] While the fluconazole resistance frequently identified among clinical isolates of *C. auris* has previously been associated with mutations in both *ERG11* and *TAC1B*, and resistance to echinocandins has been associated with mutations in *FKS1*, to date, no mechanisms contributing to clinical amphotericin B resistance have been identified in *C. auris*.[4, 6, 9-11]

We present here a case where a patient receiving treatment for a fluconazole-resistant *C. auris* infection subsequently acquired amphotericin B-resistant, and later echinocandin-resistant, disease following multiple courses of antifungal therapy. Leveraging the clinical isolates from this single case, whole genome sequencing, comprehensive sterol profiling, and Cas9-mediated genetic manipulations, we have identified the first known mechanism of clinical amphotericin B resistance in *C. auris* conferred by mutations in the sterol-methyltransferase gene, *ERG6*. Furthermore, we show that the observed mutations in *ERG6* result in abolished biosynthesis of ergosterol, the target of amphotericin B, and these mutations alone abrogate the activity of this antifungal agent.

## MATERIALS AND METHODS

### Isolate, strains, and growth media used in this study

Isolation and species-specific identification of the four clinical *C. auris* strains isolated from the patient cases included in this study was performed as described elsewhere [12]. All laboratory-derived strains and clinical isolates are listed in Supplementary Table 1 and were grown in YPD liquid medium (1% yeast extract, 2% peptone, 2% dextrose) at 35°C in a shaking incubator unless otherwise indicated. Stocks of all strains and clinical isolates were prepared with 50% sterile glycerol and were maintained at -80°C.

#### Whole genome sequencing and variant identification

Clinical isolates were cultured from glycerol stocks in YPD liquid media at 35 °C, and genomic DNA was extracted as previously described [13]. Genomic libraries were constructed and barcoded using the NEBNext Ultra DNA Library Prep kit (New England Biolabs, Ipswich, MA, USA) per manufacturer’s instructions. Genomic libraries were sequenced using the Illumina HiSeq 2500 platform with the HiSeq Rapid SBS Kit v2 as previously described [14]. Read quality and filtering was performed using FastQC v0.11.5 and PRINSEQ v0.20.3 (21278185) using “-trim_left 15 -trim_qual_left 20 -trim_qual_right 20 -min_len 100 - min_qual_mean 25 -derep 14”. Then, paired-end reads were aligned to the *C. auris* assembly strain B8441 (GenBank accession PEKT00000000.2; (30559369)) using BWA mem v0.7.12 (19451168) and variants were identified using GATK v3.7 (20644199) with the haploid mode and GATK tools (RealignerTargetCreator, IndelRealigner, HaplotypeCaller for both SNPs and indels, CombineGVCFs, GenotypeGVCFs, GatherVCFs, SelectVariants, and Variant Filtration). Sites were filtered with Variant Filtration using “QD < 2.0 || FS > 60.0 || MQ < 40.0”. Genotypes were filtered if the minimum genotype quality < 50, percent alternate allele < 0.8, or depth < 10 (https://github.com/broadinstitute/broad-fungalgroup/blob/master/scripts/SNPs/filterGatkGenotypes.py). Genomic variants were annotated, and the functional effect was predicted using SnpEff v4.3T (22728672).

#### Plasmid construction and repair template preparation

CRISPR-Cas9-mediated gene editing was performed using the transient episomal plasmid-based system described previously and the plasmid pJMR17v3 [15, 16]. Guide DNAs used as inserts in pJMR17v3 and DNA primers used in repair template generation are listed in Table 2.

#### Strain construction

Electrocompetent cells were prepared as previously described [10] and were mixed in an electroporation cuvette with ∼10 μg appropriate repair template DNA (containing the desired/introduced *ERG6* sequence) and ∼10 μg appropriate plasmid (cloned with the guide DNA matching the *ERG6* sequence of the isolate being electroporated) prior to electroporation with a BioRad GenePulser (BioRad, Hercules, CA). One milliliter 1M sorbitol was used to transfer transformation mixtures from the cuvette to a culture tube containing 1mL YPD, and transformants were allowed to recover for 4-6h at 35°C with shaking. Aliquots of the recovered transformants were spread on YPD agar plates supplemented with 200 μg/ml nourseothricin (Nou200) and incubated at 35°C until colonies formed. Single colonies were picked from transformation plates and patched sequentially on YPD agar. Once the *ERG6* sequence was verified by Sanger sequencing, desired colonies were cultured in 2mL YPD overnight at 35°C and subsequently streaked for single colonies on YPD agar. Colonies were then replica plated on both YPD agar and Nou200 to confirm nourseothricin susceptibility due loss of the pJMR17v3 plasmid.

#### Sanger sequencing

Genomic DNA was isolated from colonies with the appropriate growth phenotype and was used to amplify the *ERG6* ORF in PCR with primers CAU0013J01m and CAU0014J01m (Table 2) and Phusion Green master mix per manufacturer’s instructions (Thermo Scientific, Waltham, MA). PCR amplicons were then used as templates in Sanger sequencing reactions primed with sequencing primers (Supplementary Table 2) and run on an 3730xl DNA Analyzer (Applied Biosystem, Foster City, CA) using standard DNA sequencing chemistries.

#### Comprehensive sterol profiling

Laboratory-derived strains and the parental clinical isolates were grown to the exponential-growth phase at 35°C in RPMI liquid medium. Alcoholic KOH was used to extract nonsaponifiable lipids. A vacuum centrifuge (Heto) was used to dry samples, which were then derivatized by adding 100 μl 90% N,O-bis(trimethylsilyl)-trifluoroacetamide-10% tetramethylsilane (TMS) (Sigma, St. Louis, MO) and 200 μl anhydrous pyridine (Sigma) while heating at 80°C for 2h as previously described [10, 17]. Gas chromatography-mass spectroscopy (GC-MS) (with a Thermo 1300 gas chromatography system coupled to a Thermo ISQ mass spectrometer (Thermo Scientific) was used to analyze and identify TMS-derivatized sterols through comparison of the retention times and fragmentation spectra for known standards. Sterol profiles for each sample were determined by analyzing the integrated peak areas from GC-MS data files using Xcalibur software (Thermo Scientific). All sterol analysis was performed in biological triplicate. Error bars for each data point represent the standard deviations of results from three independent measurements of technical replicates.

#### MICs by Elipsometer test (E-test)

E-tests were performed to determine amphotericin B MICs as per manufacturer’s instructions (Biomerieux USA, Chicago, IL) with modifications as recommended by the Clinical Laboratory Standards Institute. All susceptibility testing was performed in biological duplicate.

### RESULTS

#### Patient case and antifungal susceptibility testing of clinical C. auris isolates

A 33-year old female receiving treatment for Hodgkin’s lymphoma and advanced stage nodular sclerosis was admitted to the hospital after complaints of respiratory distress. A high-resolution chest CT revealed bilateral diffuse alveolar disease, ground glass opacities, and mild right-sided pneumothorax. The patient was empirically initiated on broad spectrum antimicrobials including meropenem, linezolid, and liposomal amphotericin B (dosed at 5mg/kg) (**Figure 1**). Both bronchoalveolar lavage fluid and endotracheal tube cultures would subsequently grow *C. albicans*, and liposomal amphotericin B was continued for a total of 3 weeks. The patient’s condition improved, and chemotherapy was resumed approximately one month after the completion of amphotericin B therapy.

**Figure 1.**
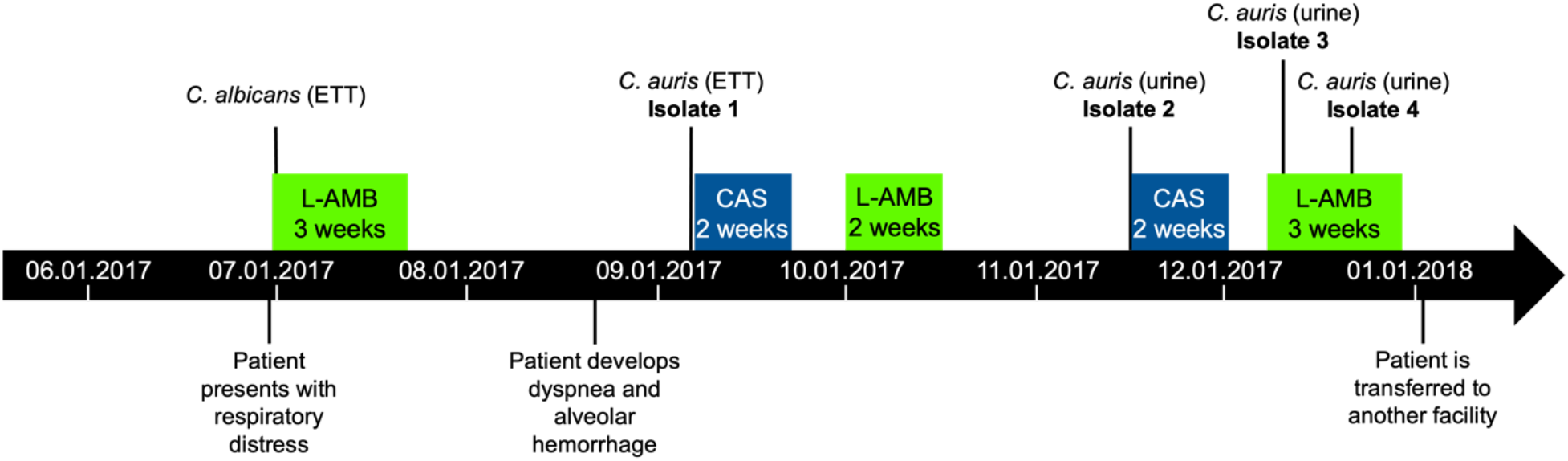
High-level amphotericin B resistance develops during treatment of a *C. auris* invasive infection. Timeline for the isolation of *Candida* clinical isolates and antifungal treatment administered to the patient. L-AMB: liposomal amphotericin B; CAS: caspofungin; ETT: endotracheal tube

Subsequently, the patient developed dyspnea and alveolar hemorrhage and treatment with meropenem and methylprednisolone was initiated. Cultures of endotracheal tube (ETT) secretions grew yeast (Isolate 1) initially identified as *Candida famata* by Vitek2, but later revealed to be *C. auris* by internal transcribed spacers (ITS) sequencing. When applying the tentative CDC *C. auris* breakpoints, Isolate 1 was resistant to fluconazole but susceptible to caspofungin and amphotericin B. A 2-week course of caspofungin treatment was initiated, to be followed by a 2-week course of liposomal amphotericin B (5mg/kg/day) when ETT cultures remained positive. Following this course of amphotericin B, all respiratory cultures remained negative for *C. auris*, however the patient remained on broad spectrum antibiotics including piperacillin-tazobactam, amikacin, and colistin for a complicated course of treatment for a carbapenem-resistant *Pseudomonas aeruginosa* respiratory tract infection.

One month later, the patient developed a urinary tract infection and cultures grew yeast initially identified as *Candida haemulonii*, but later confirmed as *C. auris* by ITS sequencing (Isolate 2). Isolate 2 was highly resistant to both fluconazole and amphotericin B but remained susceptible to caspofungin (**Table 1**), and a 2-week course of caspofungin was initiated. After the completion of caspofungin treatment, urine cultures again grew *C. auris* (Isolate 3), and a 3-week course of liposomal amphotericin B (5 mg/kg/day) was initiated. Ten days into this course of amphotericin B, urine cultures again grew *C. auris* (Isolate 4). Intriguingly, while Isolate 4 was resistant to both fluconazole and caspofungin, it had regained susceptibility to amphotericin B (**Table 1**). Shortly after the completion of amphotericin B therapy, the patient was transferred to another medical facility for further treatment and was lost to follow-up.

**Table 1.** Clinical antifungal susceptibilities for *C. auris* isolates.

| Clinical Isolate | Clinical Antifungal MIC (mg/L) |  |  |
| --- | --- | --- | --- |
|  | Amphotericin B | Fluconazole | Caspofungin |
| Isolate 1 | 0.75 | **256** | 0.75 |
| Isolate 2 | **>32** | **96** | 0.38 |
| Isolate 3 | **>32** | **96** | 0.38 |
| Isolate 4 | 0.75 | **256** | **4** |
| MIC: minimum inhibitory concentration; MIC shown in **bold** exceed tentative CDC breakpoints |  |  |  |

### Whole genome sequencing

Whole genome sequencing was performed as previously described on the four isolates and revealed all isolates to belong to Clade I (subclade b), with four or fewer single-nucleotide polymorphisms or insertions/deletions (indels) separating any two isolates, consistent with all isolates being genetically related (**Table 2**).[11] All isolates were found to have mutations previously associated with fluconazole resistance in the *ERG11* and *TAC1B* genes, encoding the Y132F and A583S amino acid substitutions, respectively. The two isolates exhibiting high-level (MIC >32mg/L) amphotericin B resistance, Isolates 2 and 3, were both found to have an indel mutation in the sterol-methyltransferase gene, *ERG6*, resulting in an early stop codon and likely nonsense transcript (YY98V*). Intriguingly, the terminal amphotericin B-susceptible isolate, Isolate 4, was found to retain this indel mutation in *ERG6*, but have also acquired a duplication of two nucleotides, resulting in a full length *ERG6* transcript with 3 altered amino acid residues (encoding RYY97LVS). Isolate 4 was also found to have a novel mutation (encoding D642Y) in hot-spot 1 of the gene encoding β-D-glucan synthase, *FKS1*.

### Impact of ERG6 mutations on Amphotericin B susceptibility

To test the influence of the identified mutations in *C. auris ERG6* on amphotericin B susceptibility, a transient Cas9-mediated *C. auris* genetic manipulation system was used to perform allelic exchange. Transformations were performed by electroporation, two independent transformants were obtained for each allele exchange, and all transformants were confirmed by Sanger sequencing as previously described [10, 11]. Introduction of the YY98V* encoding indel mutation into the *ERG6* gene of Isolates 1 and 4 resulted in a ≥32-fold increase in amphotericin B MIC in both backgrounds (**Figure 2**) and no difference was observed between biological replicate transformants (data not shown). Conversely, introduction of the wildtype (matching the antifungal susceptible B8441 reference sequence) *ERG6* sequence into both Isolates 2 and 3 resulted in a complete restoration of amphotericin susceptibility (≥32-fold reduction in MIC), confirming the role of this mutation in the high-level resistance observed.

**Figure 2.**
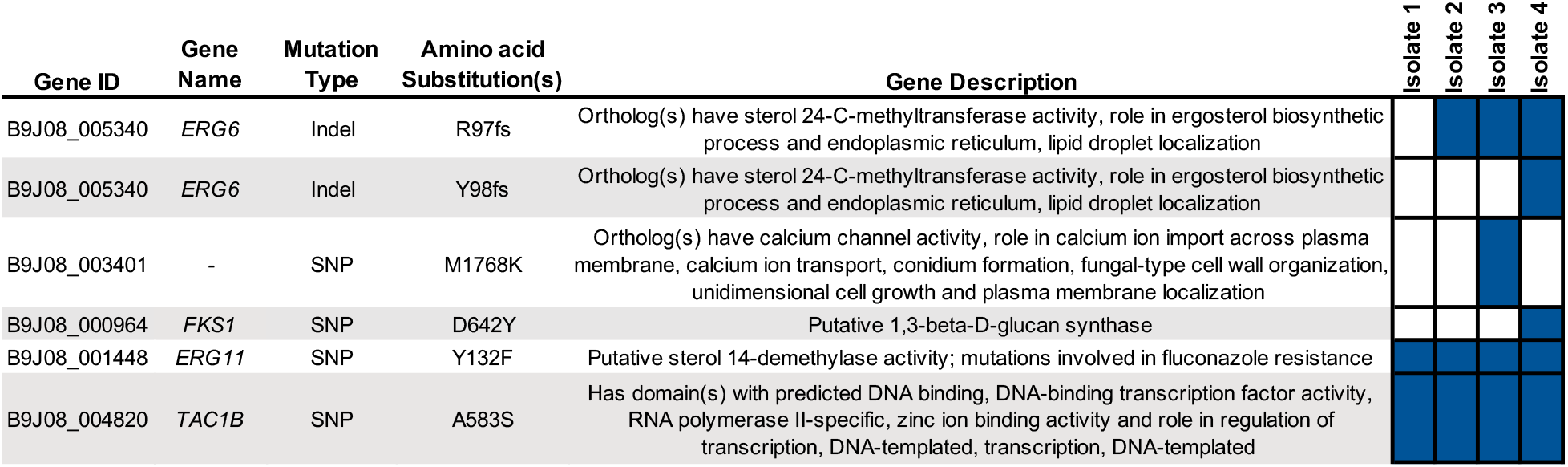
Whole genome sequencing reveals an association between mutations in *C. auris ERG6* and high-level amphotericin B resistance. Non-synonymous mutations differing between *C. auris* clinical isolates in this study, and those previously associated with antifungal resistance are shown with gene identifier, gene name, mutation type, encoded amino acid substitution(s), and gene description (as listed on the Candida Genome Database). Boxes filled in blue indicate the presence of the listed mutation in the corresponding clinical isolate.

### Comprehensive sterol profiling

As amphotericin B is known to exert an antifungal effect by directly binding to the predominant sterol of most medically relevant fungi, ergosterol, and the YY98V* encoding mutation in *ERG6* was hypothesized to result in an abrogation of sterol-methyltransferase activity, we next performed comprehensive sterol profiling to investigate changes in cellular sterol composition using methods as previously described [11]. Consistent with previously reported *C. auris* cell sterol compositions, all amphotericin B-susceptible isolates and strains were observed to have ergosterol as the predominant sterol (>60%), with ergosta-type sterols (such as ergosta-5,7-dienol and ergosta-5,8,22,24(28)-tetraenol), and early sterols (such as lanosterol and zymosterol) comprising the majority of remaining sterols (**Figure 3**). Conversely, all isolates and strains with the amphotericin B resistance-conferring YY98V* mutation in *ERG6* were found to have sterol profiles completely devoid of ergosta-type sterols. Instead, cholesta-5,7,24-trienol was found to be the most abundant sterol (>60%) in these samples, with the remainder of sterols being comprised of other cholesta-type sterols (such as cholesta-5,8,22,24-tetraenol and cholesta-7,24-dienol) and early sterols. Thus, the observed resistance to amphotericin B correlates directly with the abrogation of the production of ergosterol observed.

**Figure 3.**
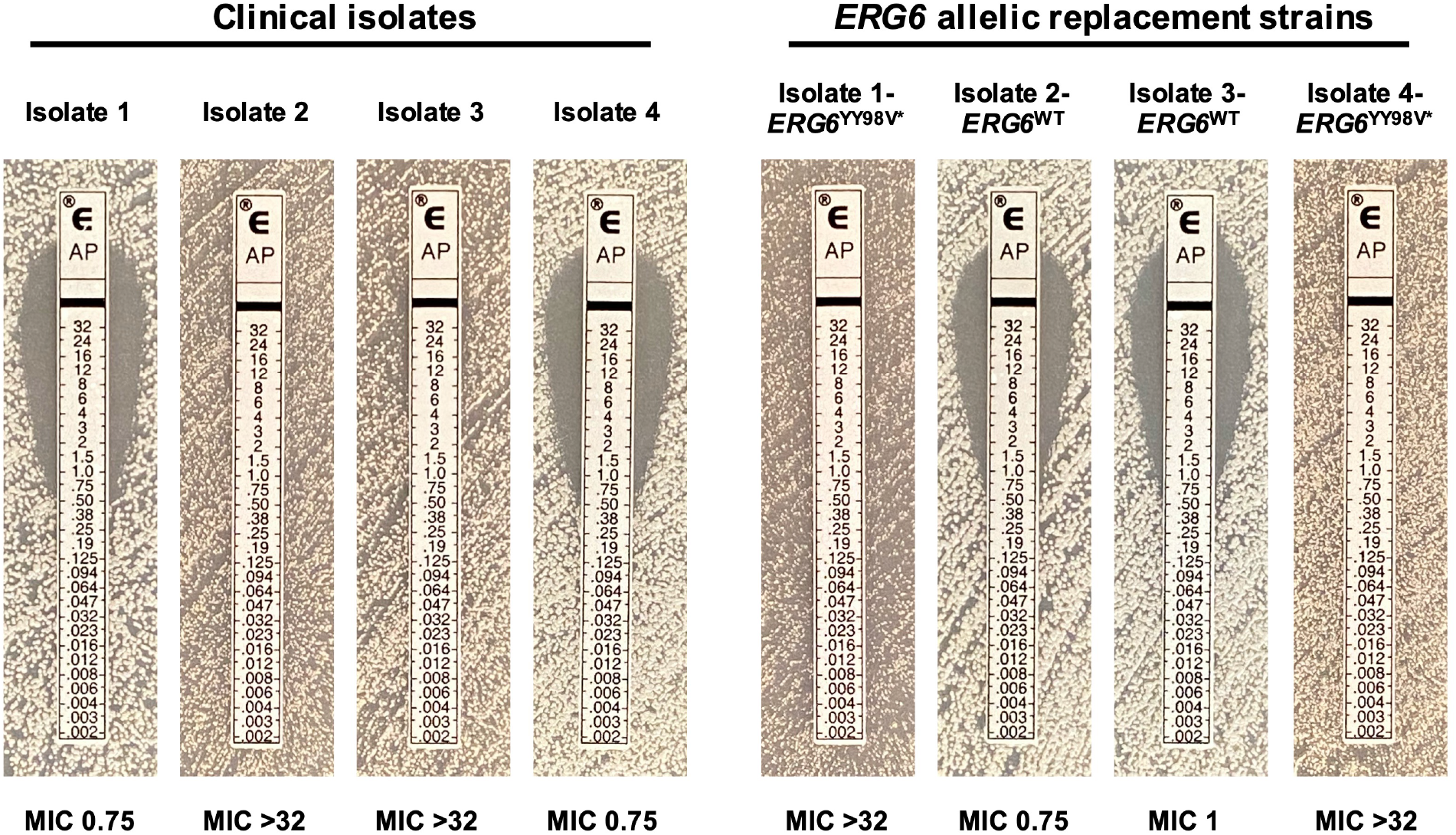
Mutations in *C. auris ERG6* confer high-level amphotericin B resistance. Amphotericin B MIC as determined by Etest for each of the clinical isolates and the *ERG6* allelic replacement strains constructed in these studies are shown.

**Figure 4.**
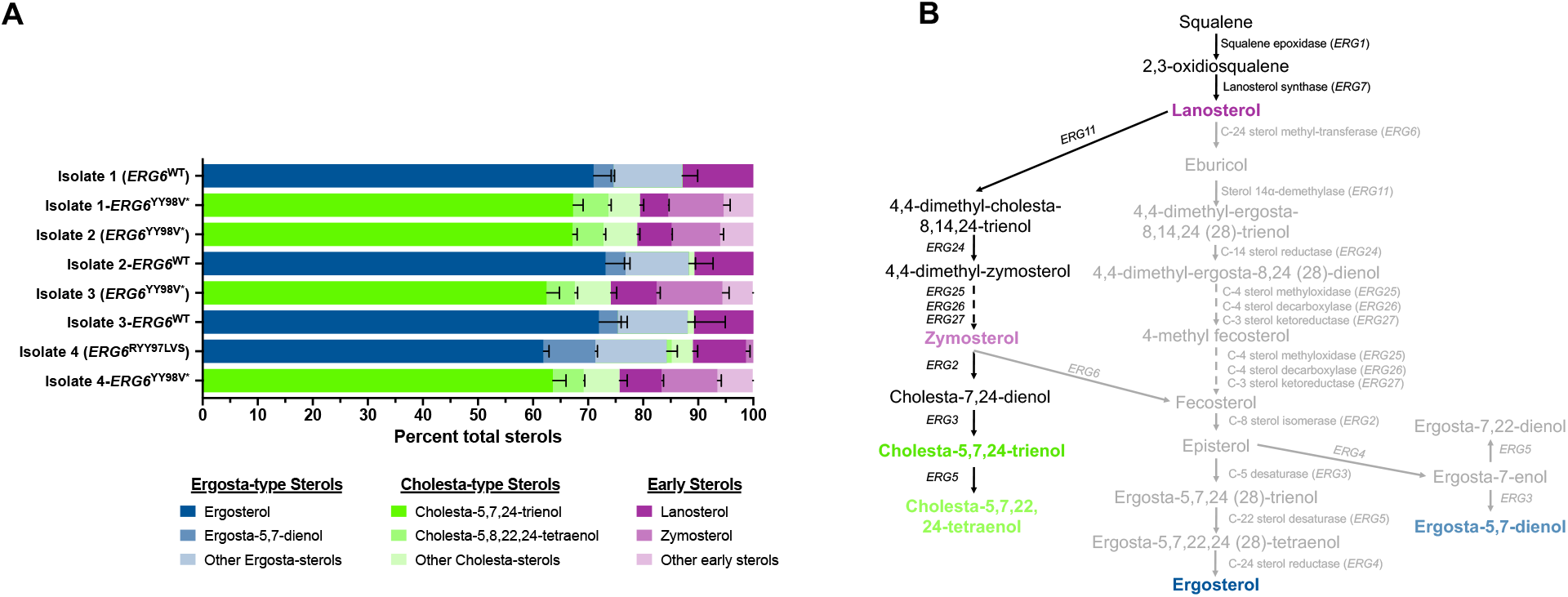
Mutations in *C. auris ERG6* result in an abrogation of ergosterol biosynthesis and accumulation of cholesta-type sterols. **A)** Comprehensive sterol profiles of each of the clinical isolates and the *ERG6* allelic replacement strains constructed in these studies. Sterol profiles are shown with each sterol represented as the proportion of total cell sterols. Error bars represent the standard deviation from three independent biological samples. **B)** Putative *C. auris* sterol biosynthesis pathway demonstrating diversion of sterol production towards cholesta-type sterols upon loss of Erg6 activity. Sterols shown in green, blue, and purple correspond to cholesta-type, ergosta-type, and early sterols as shown in comprehensive sterol profiling.

## DISCUSSION

The patient case presented here exemplifies the challenges antifungal resistance poses to clinicians treating patients with *C. auris* infections, and for the first time identifies mutations in the *C. auris* sterol-methyltransferase gene, *ERG6*, as a mechanism conferring high-level amphotericin B resistance. These findings were supported by allelic replacement studies. The *ERG6* mutations described here resulted in a dramatic shift in *C. auris* cell sterol profiles, diverting sterol production toward cholesta-type sterols more similar to those found in mammalian cell membranes. Intriguingly, while we have previously reported the combination of mutations in *Candida glabrata ERG2* and *ERG6* as being associated with both sterol profiles favoring cholesta-type sterols and increased amphotericin B resistance, the level of amphotericin B resistance observed in *C. glabrata* clinical isolates was significantly lower (MICs of 1 to 4 mg/L) than that which we observed in the *C. auris* clinical isolates in these studies (MIC >32 mg/L)[18]. It remains to be seen whether this stark difference in the magnitude of amphotericin B resistance is due to differences in other cell membrane lipids or corresponds to how these distantly related species of *Candida* respond to the stress induced by amphotericin B. Further research is needed to determine the prevalence of *ERG6* mutations among amphotericin B-resistant clinical isolates of *C. auris*.

## NOTES

### Disclaimer

The funders had no role in study design, data collection and interpretation, or the decision to submit the work for publication.

### Institutional review board statement

The clinical samples were obtained after obtaining verbal consent only as part of routine patient care and diagnostic work-up for the isolation and susceptibility testing of bacterial and fungal pathogens. The study was approved by the Health Sciences Center Ethical Committee, Kuwait University (approval letter VDR/EC/3724).

### Informed consent statement

The need for informed consent was waived by the Health Sciences Center Ethical Committee, Kuwait University as the results are reported on deidentified samples without revealing patient identity.

### Author Contributions

JMR, KSB, JFM, JEP, SLK, CAC, and PDR contributed to the conception of experimental designs and methodological development. SA, EM, AA, and RSE oversaw or contributed to the clinical and microbiological care of the patient described in this work. JMR, KSB, JFM, JEP, SA, EM, AA, and CAC performed experiments and analyzed data. JMR, SA, SLK, CAC, and PDR oversaw and the planning and execution of these studies. JMR, SLK, CAC, and PDR contributed to the financial support of this work. All authors wrote, reviewed, and approved the manuscript.

